# Differences in molecular sampling and data processing explain variation among single-cell and single-nucleus RNA-seq experiments

**DOI:** 10.1101/2022.08.01.502392

**Authors:** John T. Chamberlin, Younghee Lee, Gabor T. Marth, Aaron R. Quinlan

## Abstract

A mechanistic understanding of the biological and technical factors that impact cell and nuclear transcript measurements is essential to designing, analyzing, and interpreting single-cell and single-nucleus RNA sequencing experiments. RNA sampling in nuclei and cells is fundamentally different as nuclei contain the same pre-mRNA population as cells, yet contain a small subset of the largely-cytoplasmic mRNAs. Nonetheless, early studies argued that including pre-mRNA in single-nucleus analysis led to results comparable to cellular samples. However, typical bioinformatic workflows do not distinguish between pre-mRNA and mRNA when analyzing gene expression, and variation in the relative abundance of pre-mRNA and mRNA across cell types has received limited attention. These gaps are especially important given that incorporating pre-mRNA in routine gene expression analysis is now commonplace for both assays, despite known gene length bias in pre-mRNA capture. Here, we reanalyze public datasets from mouse and human to describe the mechanisms and contrasting effects of mRNA and pre-mRNA sampling in single-cell and nucleus RNA-seq. We disentangle the roles of bioinformatic processing, assay choice, and biological variability on measured gene expression and marker gene selection. We show that pre-mRNA levels vary considerably among cell types, which mediates the degree of gene length bias within and between assays and limits the generalizability of a recently-published normalization method intended to correct for this bias. As an alternative solution, we demonstrate the applicability of an existing post hoc gene length-based correction method developed for conventional RNA-seq gene set enrichment analysis. Finally, we show that the inclusion of pre-mRNA in bioinformatic processing can impart a larger effect on gene expression estimates than the choice of cell versus nuclear assay, which is pivotal to the effective reuse of existing data. Broadly, these analyses advance our understanding of the biological and technical factors underlying variation in single-cell and single-nucleus RNA-seq experiments to promote more informed choices in experimental design, data analysis, and data sharing and reuse.

## Introduction

Single-nucleus RNA-seq (snucRNA-seq) is typically used as a substitute for single-cell RNA-seq when dissociation of intact single cells is difficult. However, nuclei contain a minority of total cellular mRNA and are inherently enriched for pre-mRNAs **(Figure 1A)**, yielding fewer total exonic read alignments and a higher fraction of intronic read alignments(Habib et al. 2017; Hu et al. 2017). Consequently, early publications argued that lower transcript yield from nuclei could be mitigated by incorporating these intronic reads during gene expression quantification (Lake et al. 2018; Bakken et al. 2018; Lake et al. 2017; Selewa et al. 2020; Wu et al. 2019). Some studies extended this strategy to scRNA-seq to level comparisons with single-nucleus experiments (Mereu et al. 2020; Selewa et al. 2020). For the widely-used CellRanger pre-processing software, this initially required use of a custom ‘pre-mRNA’ reference annotation and later an ‘include-introns’ option, but the tool has since been updated to incorporate intronic reads by default for both cell and nuclear datasets (“Release Notes for Cell Ranger 7.0.0 (May 17, 2022): -Software -Single Cell Gene Expression -Official 10x Genomics Support “ n.d.). This paradigm shift in bioinformatic processing is important because mRNA and pre-mRNA tend to be captured by distinct sampling mechanisms with accompanying biases. Whereas mRNAs are primarily captured by the poly(A) tail in a transcript length-independent manner(Phipson, Zappia, and Oshlack 2017), pre-mRNAs are subject to a gene length-associated bias via *internal priming (Eraslan et al. 2021; Truong et al. 2022; Thrupp et al. 2020; La Manno et al. 2018; Svoboda, Frost, and Bosco 2022; “Interpreting Intronic and Antisense Reads in 10x Genomics Single Cell Gene Expression Data “ n*.*d*.*; Gorin and Pachter 2023a)*. This term refers to the hybridization of oligo(dT) primers to internal adenosine homopolymers, which frequently occur in introns and covary strongly with gene length **(Figure 1B)**. Overall gene length bias, or the degree to which the abundance of long genes is overestimated, is greater in nuclei than in cells because they contain proportionally more pre-mRNA **(Figure 1C)**.

**Figure 1:**
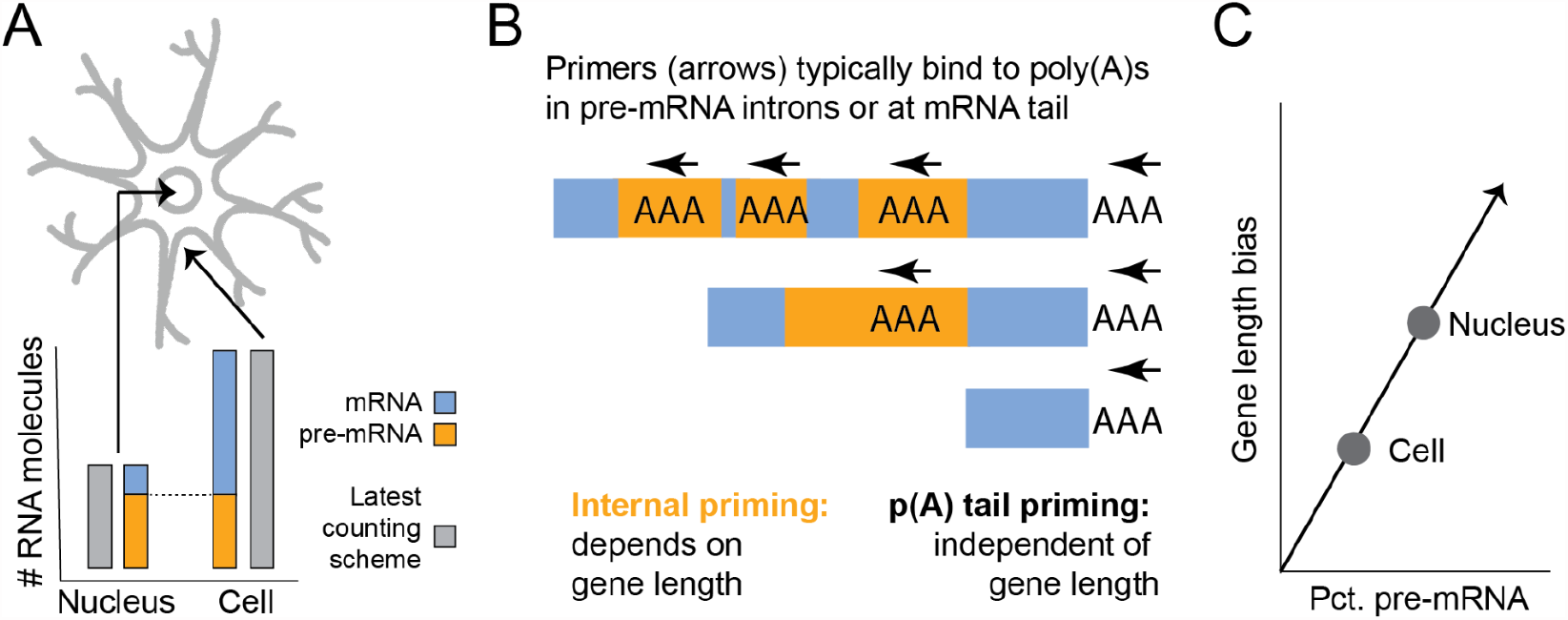
(A) Nuclei contain less total RNA but are enriched for pre-mRNA compared to cells. All pre-mRNAs in the nucleus are also in the cell, but most mRNAs are outside the nucleus (B) Pre-mRNA tends to generate intronic reads via internal priming which is biased in a gene-length associated matter. mRNA generates exonic reads from priming at the poly(A) tail, irrespective of length. Gray arrows indicate the position and direction of reverse transcription initiation by poly(dT) primers. (C) Gene length bias, or the degree to which long genes are overestimated by the assay, is a function of pre-mRNA content. Nuclei contain relatively more pre-mRNA than equivalent cells.

While a number of studies have simply noted the excess of gene length bias in snucRNA-seq (Eraslan et al. 2021; Selewa et al. 2020), others have advanced incompatible or inconsistent recommendations to account for it in different types of analyses. First, Thrupp et al argued that snucRNA-seq is fundamentally “not suitable “ as a substitute for scRNA-seq in part due to gene length bias (Thrupp et al. 2020). However, they did not comment on the role of internal priming in pre-mRNA capture, nor did they control for differences in the inclusion of intronic reads between experiments. More recently, Svodoba et al advocated for excluding internally-primed alignments to improve concordance with bulk RNA-seq(Svoboda, Frost, and Bosco 2022), while Gupta et al proposed a normalization scheme that scales down the contribution of pre-mRNA transcript counts based on gene length to improve the similarity between matched single-cell and -nucleus samples(Gupta et al. 2022). Critically, neither group demonstrated the generalizability of recommendations across cell types. This is important because pre-mRNA recovery, or implicitly, gene length bias, is known to be highly tissue-specific. For example, two early snucRNA-seq studies reported just 16% intronic read fraction in mouse heart compared to 50% in mouse brain(Hu et al. 2018, 2017). As such, the potential impact of the inclusion, exclusion, and/or normalization of pre-mRNA signals is also highly tissue-specific, and presumably cell type-specific. However, the conventional bioinformatic workflow does not separately report exonic and intronic abundances in the output gene expression matrix, so additional steps are needed to measure the level and impact of pre-mRNA across individual cells.

In summary, the presence and effect of variation in pre-mRNA sampling bias across cell types both within and between assays is obscured by standard analysis practices. Previous efforts have generally failed to disentangle the effect of assay choice (cell or nucleus) and bioinformatic processing (quantification with or without intronic reads), such that best practices in data analysis have not been established. The goal of this study is to address these gaps by reanalyzing existing experiments including a large dataset from the mouse cortex(Yao et al.2021) and the human microglia data generated by Thrupp et al. We quantify gene expression with and without including intronic alignments and reimplement the Gupta et al gene-length correction method to evaluate the effect of pre-mRNA content, and bioinformatic choices across cell types. We focus on nervous tissues because these are known to exhibit the highest levels of pre-mRNA content(Cao et al. 2020) and are frequently assayed as single nuclei, particularly in human studies. We also revisit data and metadata from an atlas of human fetal organs(Cao et al. 2020) to position our results in a broader context.

Our analysis demonstrates that observed differences are explained in part by the relative enrichment of pre-mRNAs and concomitant sampling bias, which also constrains the generalizability of the Gupta et al gene-length correction method. We quantify the necessity of utilizing identical bioinformatic processing choices to avoid technical batch effects, particularly in comparisons to pre-existing data. Finally, we recommend a modified bioinformatic workflow which allows the user to measure and anticipate the level of pre-mRNA sampling bias, and demonstrate potential post hoc solutions for improving biological interpretation of results among cell types and experiments.

## Results

### Incorporating pre-mRNA data improves similarity of cells and nuclei despite gene length bias

We focused our analysis on a well-annotated dataset from the mouse motor cortex(Yao et al. 2021). This study generated data with both SMART-seq and 10x Genomics protocols(Zheng et al. 2017) from multiple centers, not all of which included both cells and nuclei. To minimize potential batch effects, we limited our analyses to the single-cell and single-nucleus data generated by the Allen Institute using 10x Genomics V3 assay chemistry (**Methods**). We relied on the cell type annotations provided by the authors, as these were based on the integrated analysis of all datasets. We also reanalyzed human microglial data generated by Thrupp et al(Thrupp et al. 2020), also with the 10x Genomics protocol, in order to re-evaluate their conclusions with regard to bioinformatic processing choices. We aligned and quantified the sequence data from both studies with STARsolo(Kaminow, Yunusov, and Dobin 2021), a faster equivalent to the widely-used CellRanger software which allows simultaneous estimation of gene abundances with and without intronic alignments (**Methods**).

Since the nomenclature for quantification schemes is not standardized in the literature, herein we use the terms *exon* (transcript counts from exonic alignments alone), *intron* (transcript counts not from exonic alignments), and *intron&exon* (transcript counts from alignments in exons or introns). We inferred *intron* counts by subtracting *exon* counts from *intron&exon*. We also reimplemented the Gupta et al method, which we refer to as *length-corrected* counts. This entails dividing *intron* counts for each gene by the expected number of internal priming motifs in the gene (15 or more consecutive A’s) given its length, based on a genome-wide rate of .27 motifs per kilobase of gene in mouse.

To explore the impact of bioinformatic choices on estimated RNA abundances, we began with L5 IT neurons, the most common cell type in the dataset. As expected, switching both assays from *exon* quantification to *intron&exon* quantification introduced a clear gene length effect in between-assay differential expression, consistent with elevated internal priming due to pre-mRNA enrichment in nuclei (**Figure 2A,B**). Nevertheless, the net similarity between cells and nuclei actually improved, as the correlation between average gene expression increased from .74 with *exon* to .81 with *intron&exon* and the number of differentially-expressed genes decreased by about 50% (**Figure 2C,D**).

**Figure 2.**
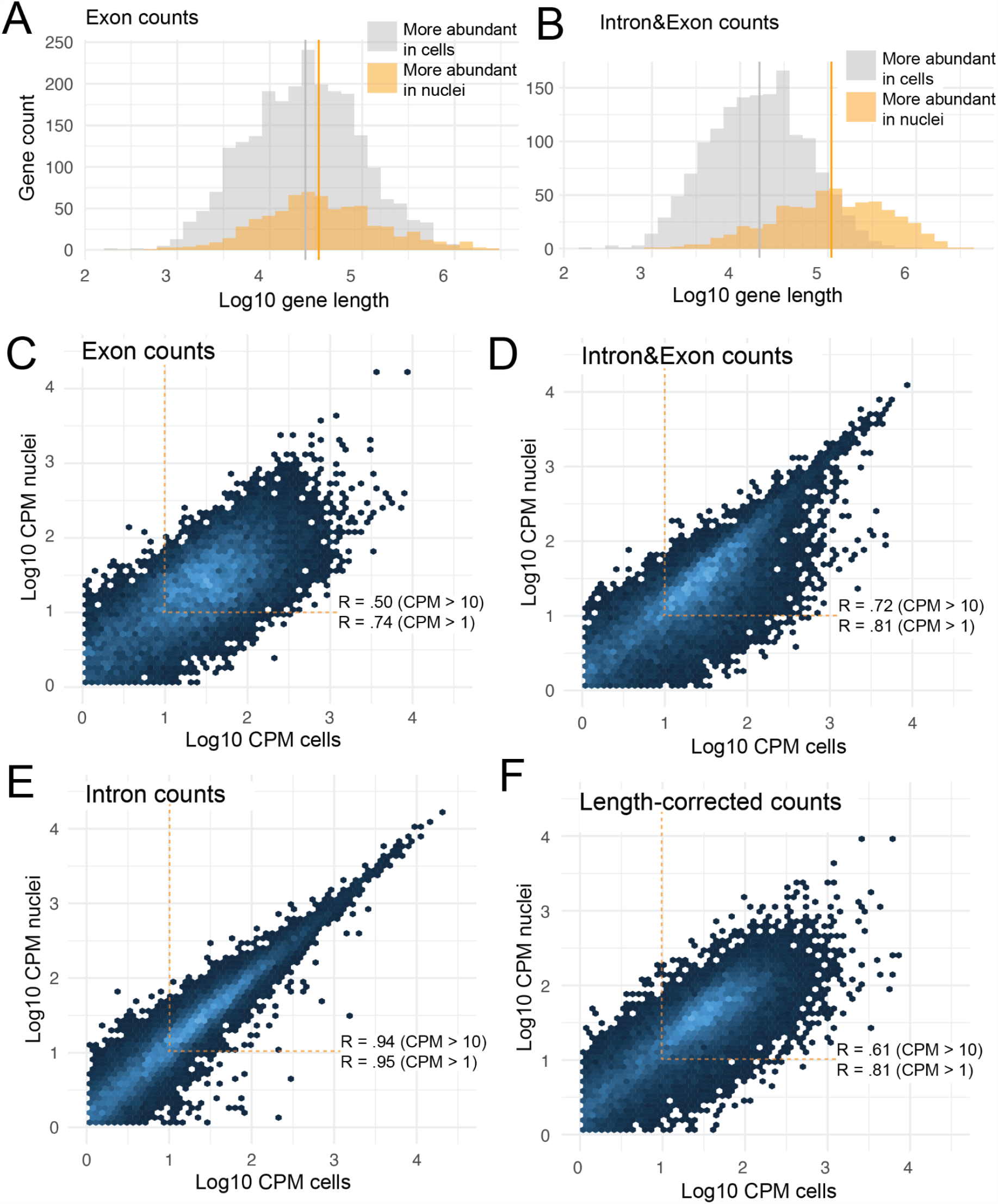
RNA sampling properties explain the effect of bioinformatic processing on assay similarity. (A) When intronic reads are excluded, genes enriched in L5 IT nuclei are only slightly longer than those enriched in cells. Mean gene lengths (Log10 scale) are 4.56 vs 4.37, p = 4.5e-12 (B) After incorporating intronic reads, the separation in gene length is much greater, but total differences decline. Mean gene lengths (Log10 scale) are 5.0 vs 4.2, p < 2e-16 (C) Correlation of *exon* abundances between L5 IT cells and nuclei. Pearson correlations are computed on log10(mean counts per million) for genes above 1 or 10 mean CPM in both assays (D) *Intron&exon* abundances are more strongly-correlated and show fewer total differences (E) *Length-corrected* abundances are no better correlated than the baseline result. Total differences increase, consistent with the worsened correlation among more highly-expressed genes (F) Correlation of *intron* abundances is very high, consistent with pre-mRNA localization within the nucleus which is within the cell. The length-correction method depresses *intron* counts which indirectly amplifies the prominence of *exon*.

In order to understand this apparent contradiction, we compared *intron* counts between cells and nuclei, which revealed a very strong correlation of .94 and just 37 differential-expressed genes (**Figure 2E)**. This indicates that pre-mRNA sampling is very similar between L5 IT neuronal nuclei and cells, while mRNA is inherently less similar. This is consistent with the subcellular localization patterns of the two RNA species: all pre-mRNAs are inside the cell, but not all mRNAs are inside the nucleus. Downsampling the *intron&exon* counts to the same depth as the *exon* counts did not change the correlation between cells and nuclei, indicating that the improvement over *exon* is not also due to a decrease in sparsity.

The Gupta et al length-correction method is intended to improve upon *intron&exon* similarity by reducing gene length bias. Gupta et al reported a ∼40% decrease in differentially-expressed genes and an increased correlation in average gene expression from 0.5 to 0.6 in a comparison of human preadipocyte cells to nuclei. However, we found the opposite effect in L5 IT neurons (**FIgure 2F)**: differential genes increased from 492 to 654 and correlation did not improve. We attribute this to large differences in pre-mRNA enrichment between preadipocytes and L5 IT neurons. In preadipocytes, the intron read fraction was both lower overall and more different between nuclei and cells (40% vs 9%, 4.4-fold). In L5 IT neurons, the levels were 66% vs 40%, an enrichment ratio of just 1.5-fold. Given that length-correction reduces pre-mRNA but not mRNA counts, this suggests that reducing gene length bias is counteracted to a varying extent by the increased emphasis on mRNA differences in a sample-specific manner.

### Variation in pre-mRNA recovery explains bias within and between assays

To understand variation in sampling bias across cell types and species, we next extended the analysis to the full mouse cortex dataset and to the human microglia data from Thrupp et al. For each cell type, we estimated pre-mRNA content as the fraction of total *intron&exon* transcript counts contributed by *intron* alignments. While we expected pre-mRNA content to vary among cell types, we were surprised to find that a higher intronic fraction in nuclei did not necessarily equate to a higher fraction in cells of the same annotated type (**Figure 3A**, p = .45). In other words, the relative enrichment of pre-mRNA between nuclei and cells varies across cell types, such that the magnitude of gene length bias between assays is unpredictable *a priori*. Using *intron&exon* counts from nuclei and cells for each cell type, we found that the count of differentially-expressed genes between assays increased with the intron content ratio between nuclei and cells (**Figure 3B**, Spearman’s ρ = .92). Human microglia, despite coming from a different species and study, clustered among the other non-neuronal cell types from the mouse dataset **(Figure 3B**, yellow diamond), suggesting that differential pre-mRNA enrichment is a general mechanism mediating measured similarity.

**Figure 3.**
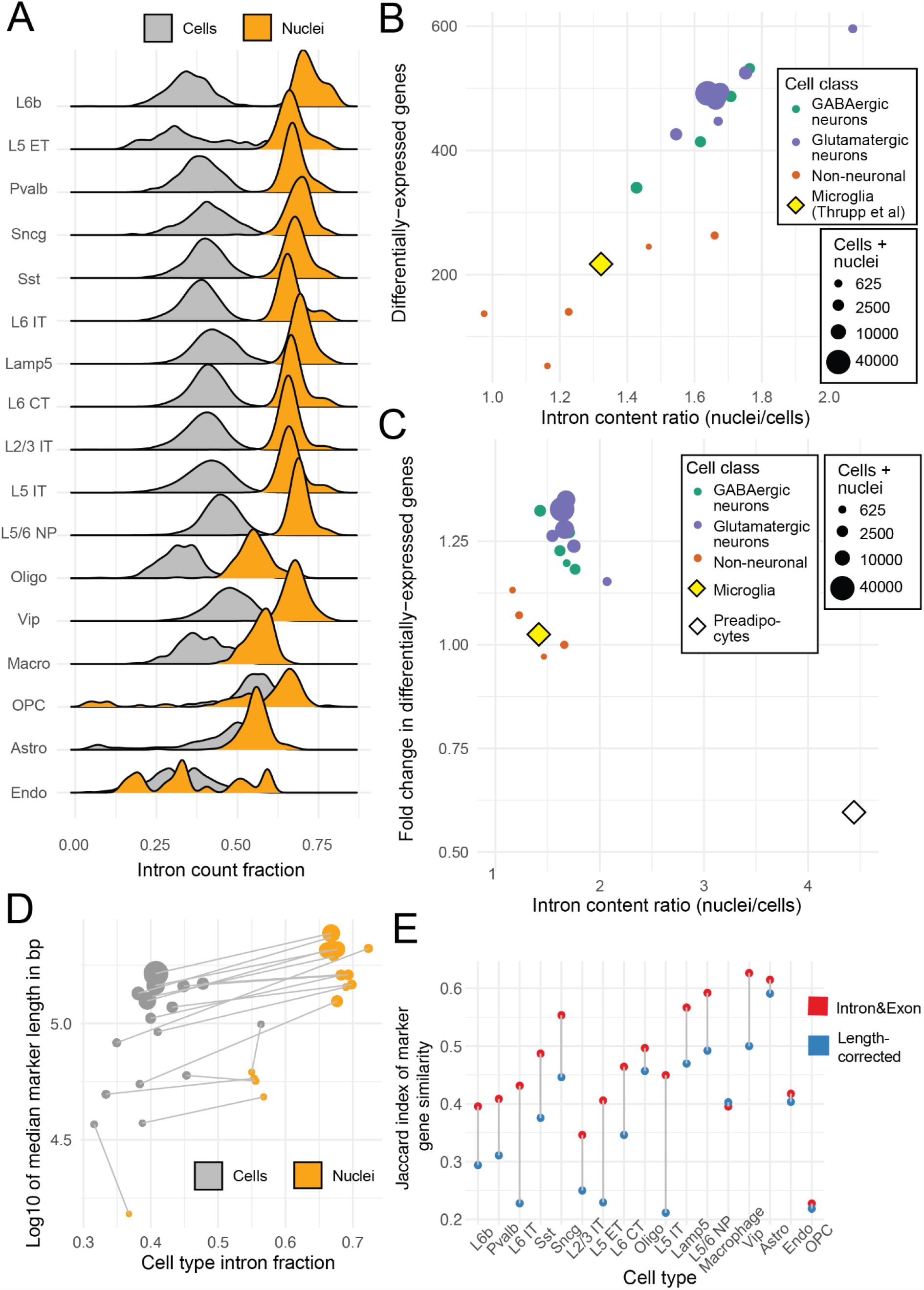
Variation in pre-mRNA enrichment moderates assay similarity with and without gene length normalization (A) Intron content distribution (total *intron* counts divided by total *intron&exon* counts per cell or nucleus) for each cell type. Mean intron content in cells does not increase with mean intron content in nuclei (Spearman’s rho = .19, p = .46). Non-neuronal cells are abbreviated: astro, astrocytes; endo, endothelial cells; oligo, oligodendrocytes; OPC, oligodendrocyte precursor cells; macro, macrophages. The remaining labels specify types of neuronal cells. (B) Differential expression of cells vs nuclei for each cell type. The number of differentially-expressed genes (log2FC > 1) increases with the ratio of mean intron content in the cell type (rho = .89, p = 9.9e-7). Cells are colored by cell type class (C) Fold change in the number of differentially abundant genes for cells vs nuclei after applying the Gupta et al length-correction procedure. The red point shows the result reported by Gupta et al for white preadipocytes. Values greater than 1 indicate adverse performance of their method. Linear modeling (of cortex cells only) identifies cell class and intron content ratio as significant predictors of method performance (R^2^ = .78, p = .0001). (D) Marker genes discovered from nuclei tend to be longer than those from cells of the same type, except for in very rare cell types. (E) Marker gene similarity (Jaccard index) for the top 50 markers (by fold change) is variable, but not in relation to the number of cells or the intron content ratio. Columns are ordered by total (cells+nuclei).

The gene length-correction method from Gupta et al was generally ineffective, showing worsened similarity for human microglia and 15 of the 17 mouse cortex cell types (**Figure 3C, Supplementary Figure 1**). We emphasize that these results are not incompatible with the values reported by Gupta et al (depicted as an open circle in **Figure 2E**), where pre-mRNA enrichment (i.e., intron content ratio) between preadipocyte nuclei and cells was much stronger than any cortex type. Instead, it suggests that the utility of their method is likely to be limited to cell types where gene length bias contributes more strongly to inter-assay differences, i.e., when pre-mRNA recovery is highly different between nuclei and cells.

While the direct comparison between assays serves as a useful benchmark, it is also important to address how these mechanisms translate to downstream analyses that would be performed in a typical, single-assay experiment. We performed marker gene analysis, which identifies genes with significantly higher expression in each cell type within a sample. Systematic gene length bias was again evident, as marker genes were consistently longer in nuclei than in cells except in the rarest cell types (**Figure 3D**). We compared the similarity of the top 50 marker genes between the two assays for each cell type using the Jaccard similarity index, or the fraction of total markers discovered by both. Unlike the previous pairwise analysis, marker similarity was not correlated with pre-mRNA enrichment; this is likely because marker gene testing is a function of all cells in aggregate. Gene length correction slightly worsened the marker gene Jaccard similarity for 15 of 17 cell types, consistent with the observed decrease in direct similarity (**Figure 3E**).

### A strategy for post hoc correction of gene length bias

Increased sampling of pre-mRNA from longer genes results in increased statistical power to detect differential expression in these genes, and vice versa. This phenomenon is similar to the fragmentation bias in conventional RNA-seq, where longer transcripts yield more fragments that can be sequenced and, in turn, more stable expression estimates, irrespective of scaling. In the context of gene set enrichment analysis, this length bias was addressed by the GOseq package published in 2010. We reasoned that GOseq could also apply to single-cell RNA-seq(Young et al. 2010, 2012). As a proof of principle, we applied the algorithm to the set of genes enriched in L5 IT nuclei compared to cells with gene length supplied as the bias term. GOseq models the relationship between the bias term (in this case, gene length) and the chance of appearing differentially expressed (**Figure 4A**). Without bias correction, the most overrepresented categories were terms indicative of neuronal function, such as “synapse organization “, which is not biologically informative given that these cells and nuclei are ostensibly the same cell type.

**Figure 4.**
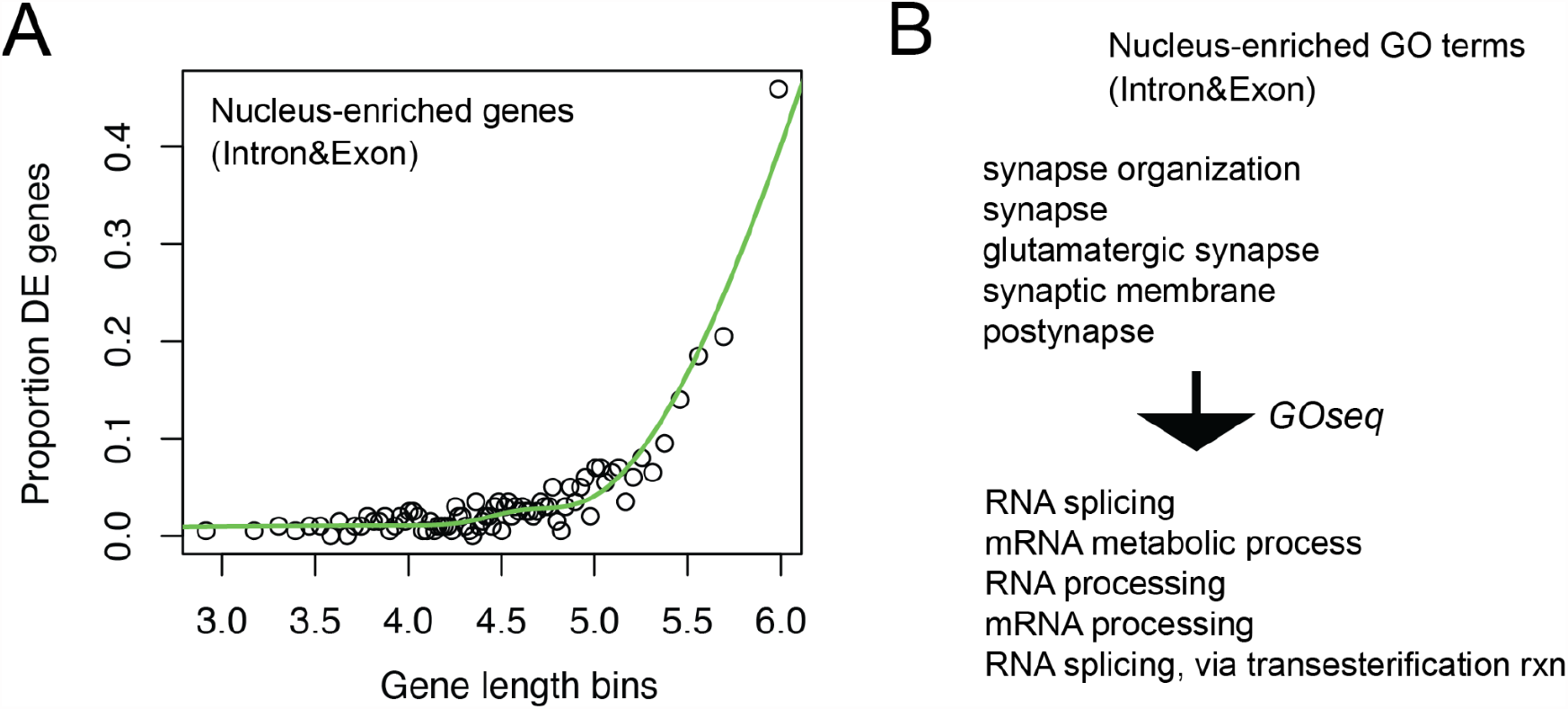
Approaches for measuring and mitigating sampling bias (A) GOseq depicts and models the relationship between a bias term (gene length) and differential expression. (B) After correcting for gene length, GO terms enriched in nuclei are consistent with known patterns of RNA localization.

After correcting for gene length, the top terms were consistent with patterns observed from RNA localization assays, such as “RNA splicing “ (**Figure 4B, Supplementary Table 1**)(Fazal et al. 2019): Fazal et al reported that “mRNAs enriched in nuclear locations tend to code for proteins enriched in nuclear speckles and nucleoplasm. “ A similar pattern was reported by Bakken et al 2018 using a technically distinct scRNA-seq assay(Bakken et al. 2018). Many of the same terms were also produced from *exon* differential expression and from the length-corrected counts, without applying GOseq (**Supplementary Table 1**). This demonstrates that gene length bias can be modeled and accounted for *post hoc* to achieve a biologically-meaningful interpretation without manipulation of raw gene counts and without discarding intronic reads, which are useful for improving cell type resolution(Bakken et al. 2018). Given that gene length bias is evident in marker genes derived from within a given single-nucleus experiment (**Figure 3D**), we suggest the adoption of GOseq when performing gene set enrichment analysis in these more commonplace scenarios as well.

### Bioinformatic processing can have a larger role than assay choice

All prior analysis performed comparisons with fixed quantification schemes, which is relevant to the current paradigm of a unified bioinformatic processing workflow. However, it does not address scenarios where data are pre-processed discordantly. This includes the historical convention in which cells were quantified as *exon* and nuclei as *intron&exon*, but will also occur if new single-cell data are compared to older, already-processed data (**Figure 5A**). Thrupp et al 2020 is a case of the former. Their study is a response to the apparent failure of human snucRNA-seq studies(Del-Aguila et al. 2019; Grubman et al. 2019; Mathys et al. 2019) to reproduce results from two earlier scRNA-seq studies in mouse models(Sala Frigerio et al. 2019; Keren-Shaul et al. 2017). These motivating studies were analyzed under their contemporary paradigms: scRNA-seq as *exon*, and snucRNA-seq as *intron&exon*. In effect, this maximizes potential differences because the results reflect both assay and bioinformatic effects. However, Thrupp et al analyzed their newly-generated data under the unified *intron&exon* method, which means they are only able to address the role of the assay.

**Figure 5.**
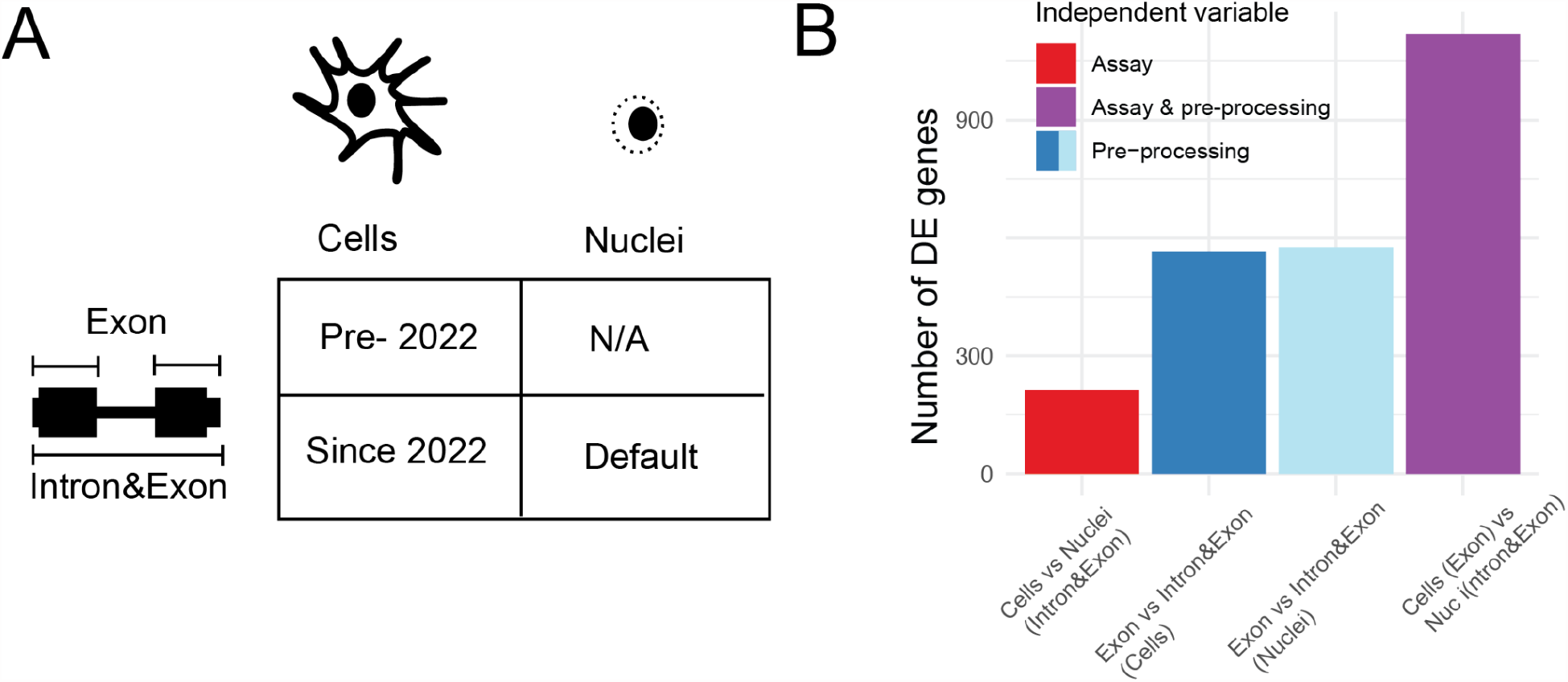
(A) Recommended bioinformatic processing of scRNA-seq and snucRNA-seq has converged over time according to CellRanger documentation (B) Processing method (blue bars) has a larger effect on differential expression than assay choice (red bar) in microglia. Historical paradigm in which nuclei and cells use two different processing methods maximizes apparent differences (purple).

We performed additional differential expression tests using Thrupp et al data. In the discordant test (cells as *exon*, nuclei as *intron&exon*), we observed 1119 differentially-expressed genes, while *intron&exon* in both assays yielded just 213 (**Figure 5B**). To understand the role of the quantification method in isolation, we also compared *exon* to *intron&exon* within cells and nuclei separately. This yielded 566 and 579 preprocessing-specific differentially expressed genes in cells and nuclei respectively. In other words, inclusion of intronic reads had a larger effect on differential gene expression than the assay itself in the Thrupp et al microglia dataset. Due to the shift in default behavior of CellRanger(“Release Notes for Cell Ranger 7.0.0 (May 17, 2022): -Software -Single Cell Gene Expression -Official 10x Genomics Support “ n.d.), the risk of repeating this type of error is likely to increase. While the incompatibility is acknowledged in principle by the software developers, it will be incumbent upon the analyst to properly account for it in practice. For example, an analysis of GTEx data by Eraslan et al also compared snucRNA-seq to scRNA-seq without controlling for differences in pre-processing, although this had little bearing on their primary analysis(Eraslan et al. 2021).

More fundamentally, our analysis demonstrates the utility of estimating per-cell/nucleus pre-mRNA content, a metric which is typically omitted by prior studies. One exception is Cao et al 2020, who included the metric in metadata but did not discuss it in their published analysis. Revisiting these data (**Supplementary Figure 2**) shows that pre-mRNA yield is generally lower in non-nervous tissues, which suggests the effect of pre-processing will be weaker, but also variable, which suggests the need to consider cell type-specific effects. As such, we implore modifying the workflow to utilize both quantification strategies. Such an informative quantification strategy is straightforward with all contemporary tools, including CellRanger(Zheng et al. 2017), alevin(Srivastava et al. 2019), kallisto(Melsted et al. 2021), STARsolo(Kaminow, Yunusov, and Dobin 2021), and simply requires two data matrices per experiment instead of one (**Methods**). Alternatively, the spliced and unspliced counts already generated by the popular Velocyto(La Manno et al. 2018) method can be substituted; nuances in these alternatives are discussed elsewhere(Soneson et al. 2021).

Simply put, estimating pre-mRNA content variability across cell types allows one to determine if observed gene length bias is attributable to the level of pre-mRNA and identify biological comparisons where *post hoc* correction is warranted, such as differential expression testing between cell types or conditions in a snucRNA-seq experiment. More generally, access to both *exon* and *intron&exon* qualifications will ensure compatibility with external datasets without the labor and computational expense of revisiting raw sequence or alignment files.

## Discussion

Single-cell sequencing technologies have advanced rapidly in the past decade alongside developments in software algorithms and bioinformatic protocols. Both factors have challenged the establishment of best practices in data analysis. In this study, we have addressed the roles of assay choice (cell or nuclear preparation) and data pre-processing method (quantification with or without intronic reads, or gene length-correction) on the consistency of results within and between datasets. Inclusion of intronic reads in snucRNA-seq expression analysis was originally borne out of a need to overcome data sparsity, but has since expanded to single-cell analysis. We confirm that universal inclusion of intronic reads can lead to a substantial improvement in direct measures of concordance between cells and nuclei, despite the introduction of gene length bias. However, the basis of this improvement is complex. In isolation, pre-mRNA measurements are more similar than mRNA measurements, reflecting patterns of synthesis, localization, and degradation within the cell. Differential enrichment of pre-mRNAs between cells and nuclei contributes to remaining differences via gene length bias. The bias also manifests in results from within a given experiment, as cell type-defining marker genes derived from snucRNA-seq will be systematically longer than from equivalent scRNA-seq. As such, pre-mRNA has a nuanced impact on inter-assay similarity: it introduces gene length bias, but also buffers against inherent noise in mRNA sampling. More fundamentally, these results show that bioinformatic choices depend on the specific aims of the analysis. For example, previous analyses have shown that incorporating intronic reads does not improve similarity to bulk RNA-seq(Del-Aguila et al. 2019), which suggests that the procedure may not be helpful for applications such as bulk sample deconvolution. Similarly, the high consistency of pre-mRNA measurements implies that the best way to minimize apparent differences between assays would be to exclude mRNA data entirely. This is obviously an untenable recommendation and suggests that orthogonal approaches are needed for evaluating pre-processing methods.

Collectively, these findings lead us to a series of recommendations. Foremost, the quantification method must be controlled for. Newly-generated data are likely to be incompatible with previously-published datasets unless care is taken to harmonize the bioinformatic workflow, and the same is true for independent cell and nuclear datasets. Fortunately, modifying the bioinformatic workflow to quantify gene expression both with and without introns is straightforward to implement and has further utility for measuring the levels of pre-mRNA sampling bias across cell types. Separating *exon* and *intron* counts has also been used for other steps in single-cell analysis, such as ambient RNA decontamination(Alvarez et al. 2020) and in mechanistic models of differential expression(Gorin and Pachter 2023b).

While the high absolute levels of pre-mRNA in nervous tissue cells and nuclei appear to preclude the use of the Gupta et al length-correction method in its current state, performance in other tissues is likely to be positive. Conversely, selected tissues such as muscle and heart tend to yield very little pre-mRNA, leading to a low level of gene length bias which may not warrant correction. However, the variable pre-mRNA enrichment evident across cortex cell types indicates that the length-correction method will require validation with matched single-cell and nucleus datasets before employing it in new contexts.

The high consistency in pre-mRNA sampling helps to reduce apparent differences between assays, but it does not ameliorate gene length bias within a given assay. Specifically, marker genes from nuclei or high pre-mRNA tissue types will be artifactually longer than markers from low pre-mRNA samples. The same logic applied to findings from within a given experiment or between conditions. Increased power for measuring longer genes, and the depletion of measurements from short genes, is an inherent feature of the assay, irrespective of scaling. As such, we suggest use of GOseq to measure and correct for this bias when performing gene set enrichment analysis.

Our analysis is subject to a number of limitations. Foremost, we relied on the mouse cortex cell type annotations generated by Yao et al, as these were based on the integrated analysis of seven datasets. Further effort is needed to assess the impact of pre-mRNA content levels on cell type assignment. However, clustering and cell type assignment have recently been reported to be robust to the effects of ambient RNA contamination, a distinct technical challenge in the field(Janssen et al. 2023).

Additional limitations are relevant to the Gupta et al method. We did not alter the parameterization of their approach. In particular, the definition of an internal priming site is specified arbitrarily, but it directly determines the reduction in total *intron* counts. Currently, *intron* counts are divided by the expected number of A(15) motifs of the gene, given its length. Relaxing the motif definition (e.g., A(10) or allowing mismatches) would result in a substantially greater reduction in total counts, and vice versa. *Exon* and *intron* quantification also do not perfectly correspond to pre-mRNA and mRNA or their capture modes(Gorin and Pachter 2023b). Others have reported that gene length bias is not fully explainable by internal priming sites alone, and is also present in *exon* counts to a lesser extent(Kuo, Hansen, and Hicks 2022). True biological differences, or additional technical artifacts(“Interpreting Intronic and Antisense Reads in 10x Genomics Single Cell Gene Expression Data “ n.d.) may also contribute. We speculate that the Gupta et al length-correction method could be improved by instead dividing counts into poly(A) tail- and internally-primed, deriving the scaling factor empirically, and flooring the scalar at 1 to avoid variance-inflation of short genes. Beyond computational solutions, these issues would likely be substantially diminished through improved single-nucleus protocols which retain more nuclear-associated mRNAs during dissociation(Drokhlyansky et al. 2020).

The description of gene length bias in single-cell and nucleus sequencing is not a novel report, but solutions remain almost absent in practice. Meanwhile, discussions have tended to emphasize open questions in so-called ‘downstream’ analysis steps of single-cell data science(Lähnemann et al. 2020; Heumos et al. 2023). However, upstream and downstream analyses are clearly not independent tasks, and further effort and attention is needed to establish unifying principles of data analysis and interpretation as the field continues its rapid expansion(Svensson et al. 2020)

## Methods

### Single-cell/nucleus RNA-seq data pre-processing

We analyzed the 10x Genomics V3 data subset from Yao et al(Yao et al. 2021) generated at the Allen Institute for Brain Studies; a complementary dataset generated at the Broad Institute was excluded because it included nuclei only. Raw sequence data and metadata were downloaded from http://data.nemoarchive.org/biccn/lab/zeng/transcriptome/. Thrupp et al data were acquired from GEO accessions GSE153807 and GSE137444.

Alignment and gene expression quantification were performed using STARsolo(Kaminow, Yunusov, and Dobin 2021) version 2.7.3a with option “--soloFeatures Gene GeneFull “, which corresponds to the *exon* and *intron&exon* schemes. We used the GRCm38 and GRCh38 reference genomes for the mouse and human samples, respectively, and Ensembl version 99 gene annotations filtered by biotype based on 10x Genomics guidelines and the “cellranger mkgtf″ utility. Due to erroneous resequencing of some original mouse cortex libraries, we used seqtk(Li n.d.) to filter out read pairs with truncated barcodes (shorter than the intended 28 base pairs), as the mixture of barcode read lengths was incompatible with STARsolo. Resulting abundance estimates were nearly identical to CellRanger values provided by the authors, which is expected by design.

Microglia snucRNA-seq data contained a mixture of cell types but barcode annotations were not provided, so we replicated the Seurat-based clustering analysis as closely as possible based on the description in Thrupp et al. We identified two clusters that corresponded to microglia based on marker genes described by Thrupp et al, including P2RY12. The single-cell data contain only microglia as they were FACS purified before sequencing, obviating the need for cluster analysis.

Preprocessed data and metadata from the human fetal atlas study(Cao et al. 2020) were downloaded from NCBI GEO accession GSE156793.

### Statistical analyses

Primary analyses were conducted in Rstudio using Seurat(Butler et al. 2018) and tidyverse packages(“Welcome to the Tidyverse “ n.d.). Differential expression tests were performed with the FindAllMarkers function. For marker gene analysis, cell types were used as identity classes. To test differences between assay- and/or pre-processing, the relevant Seurat objects were merged as needed. We used a fold change threshold of 1.5 and tested for upregulated markers only. The number of A(15) motifs in the mouse genome was calculated with Biostrings(Pages et al., n.d.) and GenomicRanges(Lawrence et al. 2013) R packages.

### Estimation of pre-mRNA content and scaling approach

We inferred pre-mRNA content from the difference in total transcripts detected with or without the use of intronic reads. STARsolo refers to these approaches as “Gene “ (Exon) and “GeneFull “ (Intron&Exon). We filtered the ‘raw’ matrices to include only the cell barcodes defined by Yao et al. We re-implemented the Gupta et al length-correction approach as described in their paper and code. *Intron* counts were taken as the difference between *Intron&Exon* and *Exon*, floored at 0. *Intron* counts were then divided by a gene-specific scalar, the expected number of A(15) motifs per gene, given gene length, and added back to *exon*. We calculated a scalar of .27 motifs per kilobase of gene in mouse.

### GOseq analysis

Genes significantly more abundant in L5 IT nuclei compared to cells (fold change > 1.5) were used as input for GOseq; for the background set, we used genes that were detected with at least 1 CPM in both cells and nuclei. The same approach was used for each pre-processing mode. We supplied gene length as the bias term, as opposed to transcript length used by the developers.

All analysis code is publicly available at https://github.com/johnchamberlin/internalpriming

## Acknowledgements

JC was supported by an NLM T15 training grant in biomedical informatics, project number 5T15LM007124-25.

## Supplementary Materials

**Figure S1:**
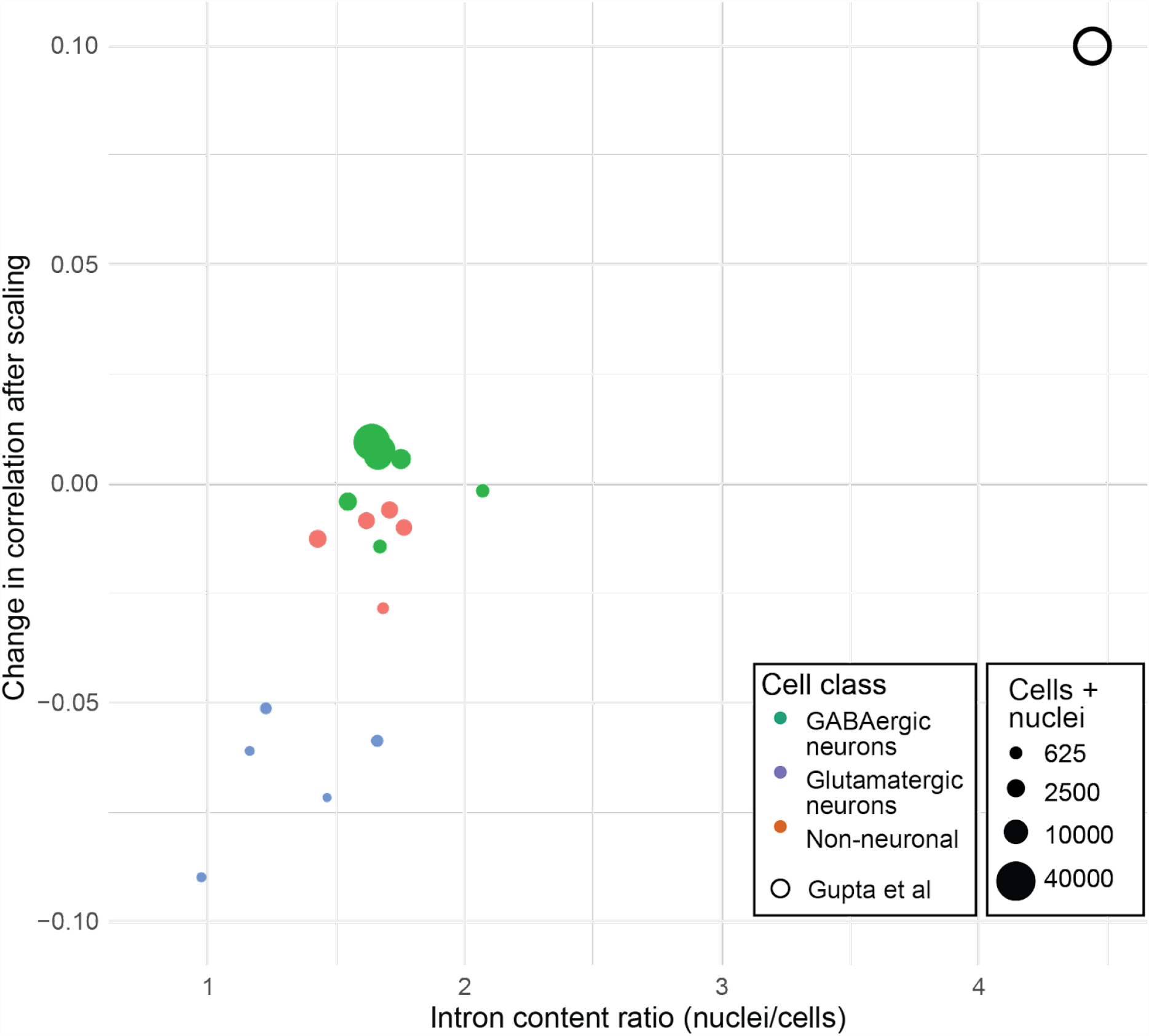
Effect on normalization on correlation between cell and nuclear abundances in mouse motor cortex. Points are colored by cell category and sized by the total number in both assays. Result reported by Gupta et al is shown as an open circle.

**Figure S2:**
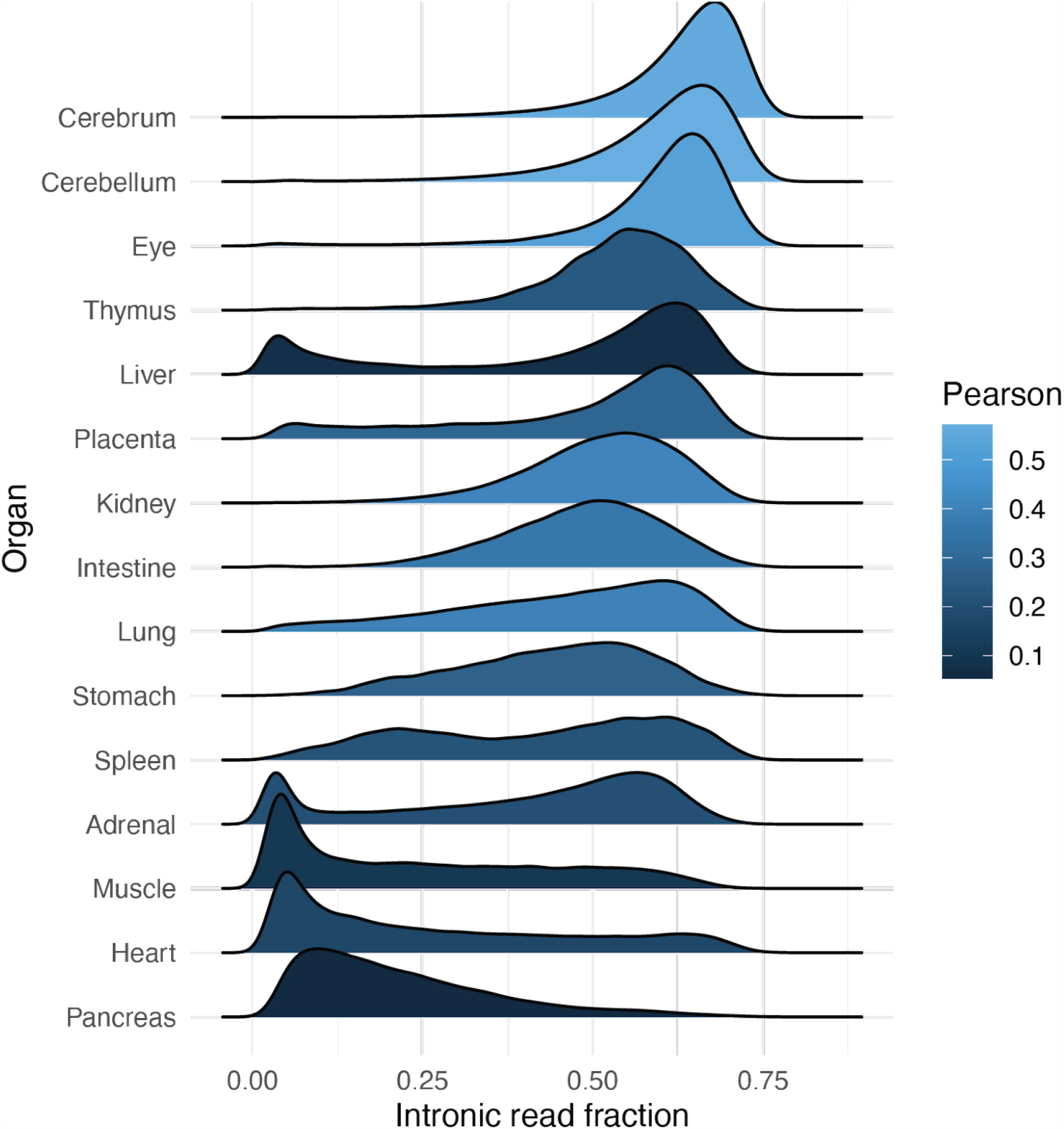
Variation in intronic read fraction reported as metadata by the Human Fetal Atlas(Cao et al. 2020). Tissues are ordered by median fraction; nervous tissues exhibit the strongest correlations between average *intron&exon* abundance (log of average CPM) and gene length. Some tissues show stark bimodality, such as Liver and Adrenal gland.

